# In-vivo Protection from SARS-CoV-2 infection by ATN-161 in k18-hACE2 transgenic mice

**DOI:** 10.1101/2021.05.08.443275

**Authors:** Narayanappa Amruta, Elizabeth B. Engler-Chiurazzi, Isabel C. Murray-Brown, Timothy E. Gressett, Ifechukwude J. Biose, Wesley H. Chastain, Gregory Bix

## Abstract

Severe acute respiratory syndrome coronavirus 2 (SARS-CoV-2) is an infectious disease that has spread worldwide. Current treatments are limited in both availability and efficacy, such that improving our understanding of the factors that facilitate infection is urgently needed to more effectively treat infected individuals and to curb the pandemic. We and others have previously demonstrated the significance of interactions between the SARS-CoV-2 spike protein, integrin α5β1, and human ACE2 to facilitate viral entry into host cells *in vitro*. We previously found that inhibition of integrin α5β1 by the clinically validated small peptide ATN-161 inhibits these spike protein interactions and cell infection *in vitro*. In continuation with our previous findings, here we have further evaluated the therapeutic potential of ATN-161 on SARS-CoV-2 infection in k18-hACE2 transgenic (SARS-CoV-2 susceptible) mice *in vivo*. We discovered that treatment with single- or repeated intravenous doses of ATN-161 (1 mg/kg) within 48 hours after intranasal inoculation with SARS-CoV-2 lead to a reduction of lung viral load, viral immunofluorescence and improved lung histology in a majority of mice 72 hours post-infection. Furthermore, ATN-161 reduced SARS-CoV-2-induced increased expression of lung integrin α5 and αv (an α5-related integrin that has also been implicated in SARS-CoV-2 interactions) as well as the C–X–C motif chemokine ligand 10 (*Cxcl10*), further supporting the potential involvement of these integrins, and the anti-inflammatory potential of ATN-161, respectively, in SARS-CoV-2 infection. To the best of our knowledge, this is the first study demonstrating the potential therapeutic efficacy of targeting integrin α5β1 in SARS-CoV-2 infection *in vivo* and supports the development of ATN-161 as a novel SARS-CoV-2 therapy.

## Introduction

Severe acute respiratory syndrome coronavirus 2 (SARS-CoV-2), the agent causative of coronavirus disease 2019 (COVID-19), is a highly transmissible respiratory pathogen in which an estimated 14% of all patients will develop serious conditions, with a subsequent mortality rate of 1.4 - 3.4% [1-3]. Several non-pharmacological interventions have been implemented to slow down the spread of SARS-CoV-2[4-7]. However, to date there is no universally agreed direct therapy available to treat COVID-19. It is important to study effective therapeutic targets focusing on disrupting aspects of the viral replication process, including SARS-CoV-2 host cell entry [8], which uses its spike proteins to bind to the angiotensin-converting enzyme 2 (hACE2) receptor on targeted cells to facilitate its entry and replication [9]. The virus infects the cells after the proteolytic cleavage of the spike protein by the transmembrane serine protease 2 (TMPRSS2) or Cathepsin B or L or FURIN is required for spike protein priming and virus infection [10].

Integrins are family of cell adhesion receptors that may play an important role in SARS-CoV-2 host cell entry and infection due to the spike protein containing an integrin binding motif arginine-glycine-aspartate (RGD) sequence [11-14]. Integrins are composed of non-covalently linked α and β subunits that recognize and bind to extracellular matrix (ECM) proteins and mediate cell survival, proliferation, differentiation, and migration [15-17]. Integrin dimers are expressed in most cells, including endothelial cells, and epithelial cells in the respiratory tract [18], and are known to be involved in the infectious etiology of other viruses such as human cytomegalovirus [19], Epstein–Barr virus [20], rotavirus [21], Kaposi’s sarcoma-associated virus (HHV-8) [22] and Ebola [23]. Importantly, the β1 family of integrins are closely associated with ACE2 [24]. A recent study reviewed novel mutation (K403R) in the spike protein that does not exist in other strains of the coronavirus, creating an RGD motif, which could be recognized by integrins [2]. Therefore, the new RGD motif in SARS-CoV-2 could increase the binding potency of ACE2-positive target cells in association with β1 integrins as well as potentially facilitating infection of ACE2-negative cells [2]. This could help explain the faster and aggressive spread of virus as compared to SARS-CoV-1, which belongs to the same family of betacoronaviruses [25]. Targeting therapies to disrupt the spike-binding event may also inhibit viral replication by halting host cell entry and offers a promising approach to treat COVID-19 [8].

Our laboratory has previously shown that the SARS-CoV-2 spike protein is capable of binding to α5β1 integrin in cell-free ELISA assays [26], an observation which has since been repeated in epithelial cell-containing systems *in vitro* [27]. We further demonstrated that the small pentapeptide α5β1 integrin inhibitor ATN-161 could inhibit this binding and also decrease SARS-CoV-2 infection and cytopathy in cultured vero-E6 cells [26]. ATN-161 has several properties that make it potentially attractive as a novel SARS-CoV-2 therapy; it is safe and well-tolerated with no dose-limiting toxicity in phase I cancer clinical trials [28], has demonstrated *in-vivo* efficacy in mice against a different beta coronavirus, porcine hemagglutinating encephalomyelitis virus (PHEV) [29] as well as other disease/injury models[30, 31], and can also bind to and inhibit integrin αvβ3 which is similar to α5β1 integrin that has also been implicated in SARS-CoV-2 infection [32, 33]. In the present study we examined the therapeutic potential of ATN-161 in a human ACE2 receptor-expressing K18 mouse model of SARS-CoV-2 infection.

## Methods

### Mice and Ethics Statement

Male heterozygous K18-hACE c57BL/6J mice (strain: 2B6.Cg-Tg(K18-ACE2)2Prlmn/J, 10-weeks-old) were obtained from The Jackson Laboratory. Mice were housed in the animal facility at Tulane University School of Medicine. The Institutional Animal Care and Use Committee of Tulane University reviewed and approved all procedures for sample handling, inactivation, and removal from a BSL3 containment (permit number 3430 (#5)).

### SARS-CoV-2 Infection

Mice were inoculated with either saline or SARS-CoV-2 via intranasal administration by the ABSL3-trained staff with a dose of 2×10^5^ TCID_50_/mouse to induce viral infection in these animals [34, 35]. The infected mice were observed daily to record body weight and clinical signs of illness (e.g. fur ruffling, less activity). After 3 days post infection (dpi), the mice were euthanized by CO2 asphyxiation followed by cervical dislocation and lungs were harvested for histology, immunofluorescence and qRT-PCR analysis.

### Treatment Interventions

Mice received ATN-161 (1 mg/kg, Medkoo Biosciences, Morrisville, NC, USA) via retro orbital i.v. injections. The subjects were divided into 6 groups; two control groups were non-infected: 1. saline treated control (saline, n=3) and 2. ATN-161 treated mice (ATN-161, n=3). These mice resided in the BSL1 portion of the Tulane Department of Comparative Medicine animal housing facility. Four SARS-CoV-2 infected groups received either saline or ATN-161 prepared fresh daily (3. SARS-CoV-2 + saline (n=5), 4. SARS-CoV-2 + ATN-161 once at 2 h (n=5), 5. SARS-CoV-2 + ATN-161 daily (2, 24, and 48 h, (n=5)) and 6. SARS-CoV-2 + ATN-161 once at 48 h (n=5) treatment administration following intra nasal viral inoculation with SARS-CoV-2.

### Viral copy number determination

Tissues were weighed and homogenized in Trizol Lysis Reagent (Invitrogen). RNA was extracted from lung homogenates according to the instructions of the RNA extraction kit manufacturer (RNeasy Plus Mini Kit; Qiagen) post phase separation using Trizol reagent. Total RNA (Five microliters) was added in duplicate to a 0.2-ml standard 96-well optical microtiter plate (Cat. No.N8010560; Thermo Fisher). qRT-PCR reaction was set by using TaqPath 1-Step Master Mix (Cat. No. A28527; Thermo Fisher) with a primers and a FAM-labeled probe targeting the N1 amplicon of N gene (2019-nCoV RUO Kit, Cat. No. 10006713; IDT-DNA) of SARS-CoV-2 (https://www.ncbi.nlm.nih.gov/nuccore; accession number MN908947), following the manufacturer’s instructions. Viral load was calculated by the linear regression function by Cq values acquired from 2019 nCoV qRT-PCR Probe Assays. The viral copy numbers from the lung samples are represented as copies/100 ng of RNA as followed using a published assay [36, 34, 37]. Subgenomic mRNA (sgmRNA) encoding the E gene was quantified using FAM-labeled primers (sgm-N-for: 5’-CGATCTCTTGTAGATCTGTTCTC-3’,sgm-N-Probe:5’-FAMTAACCAGAATGGAGAACGCAGTGGG-TAMRA-3’,sgm-N-reverse:5’-GGTGAACCAAGACGCAGTAT-3’), following the manufacturer’s instructions. Subgenomic N viral copy number was calculated by standard Cq values. The viral copy numbers from the lung samples are represented as copies/100ng of RNA followed using a published assay [37, 34].

### Gene expression

The lung tissue was homogenized in Trizol Lysis Reagent (Invitrogen). RNA was extracted from lung homogenates according to the instructions of the RNA extraction kit manufacturer (RNeasy Plus Mini Kit; Qiagen) post phase separation using Trizol reagent. RNA was converted to cDNA using iScript reverse transcriptase master mix (Bio-Rad). qPCR was carried out with QuantStudio□3 Real-Time PCR Systems (Life Technologies) using TaqMan PCR Master Mix and premixed primers/probe sets (Thermo Fisher Scientific) sets specific for *Itgav* (Mm00434486_m1), *Cxcl10* (Mm00445235_m1), *Itga5* (Mm00439797_m1), *Itgb3* (Mm00443980_m1), *Itgb1* (Mm01253233_m1), *ACE2* (Hs01085333_m1), and Hypoxanthine phosphoribosyltransferase, *Hprt* (Mm01545399_m1, Control gene) (Life Technologies) were used gene expression. Data were analyzed comparing control to SARS-CoV-2 infected mice and are presented as a fold change of control.

### Histology

The harvested whole right lungs (three lobes) were fixed in Z-fix (Anatech Ltd, Battle Creek, MI, USA). Paraffin sections (5 μm in thickness) were used for haematoxylin and eosin and Masson’s trichrome stains to identify morphological changes in lungs. Slides were imaged with a digital slide scanner (Zeiss Axio Scan; Zeiss, White Plains, NY). Representative photo micrographs at 20× magnification were acquired from the whole scanned right lung using the Aperio Image Scope software (version 12.3.2.8013, Leica, Buffalo Grove, IL, USA).

### SARS-CoV-2 immunofluorescence

Fixed tissue samples were processed via indirect immunofluorescence assays (IFA) for the detection of the SARS-CoV-2 antigen. The slides were deparaffinized in xylenes and rehydrated through an ethanol series, followed by heat-induced antigen retrieval with high pH antigen unmasking EDTA solution. The slides were washed with PBS with 0.1% Triton X-100 and blocked with 10% normal goat serum at room temperaturefor 1 h.Primary antibody (Polyclonal Anti-SARS Coronavirus (antiserum, Guinea Pig) 1:1000, NR-10361) incubation was achieved at room temperature for 1 h. Slides were washed and primary antibody detected following 40 min incubation in an appropriate secondary antibody tagged with Alexa Fluor fluorochromes (1:1000) in normal goat serum. After washing in PBS, mounting media with DAPI was used to label the nuclei. Slides were imaged with a digital slide scanner (Zeiss Axio Scan.Z1; Zeiss, White Plains, NY). Fluorescent images were acquired using HALO (Indica Labs, v2.3.2089.70). Threshold and multiplex analyses were performed with HALO for quantitation.

### Statistics

Statistical tests were performed with GraphPad Prism, 8.4.3 version (GraphPad Software, San Diego, CA). Data are presented as mean ± SEM. Significant differences were designated using omnibus one-way ANOVA and, when significant, followed-up with two-group planned comparisons selected *a priori* to probe specific hypothesis-driven questions (saline vs SARS-CoV-2+vehicle; SARS-CoV-2+vehicle vs each of the various SARS-CoV-2+ATN-161 treated groups); the Holm-Sidak adjustment was applied to control for multiple comparisons. Non-parametric tests were employed in the event of violations of underlying ANOVA assumptions. Statistical significance was taken at the *p* < 0.05 level.

## Results

### ATN-161 impacts the Genomic-N and Sub genomic N (Sgm-N) viral load in SARS-CoV-2 infected K18-hACE2 mice lung tissue

Lung tissue expression of hACE2 and viral loads after 3dpi with SARS-CoV-2 + Saline or ATN-161 administered either once at 2 h or 48 h post-infection or daily, post-infection, were measured, analyzed and compared to Vehicle-treated/uninfected (no SARS-CoV-2 exposure) mice either given Saline or ATN-161 (daily) for 3dpi. (Fig 1A). For hACE2, the omnibus one-way ANOVA was not significant (Fig 1B).

**Figure 1.**
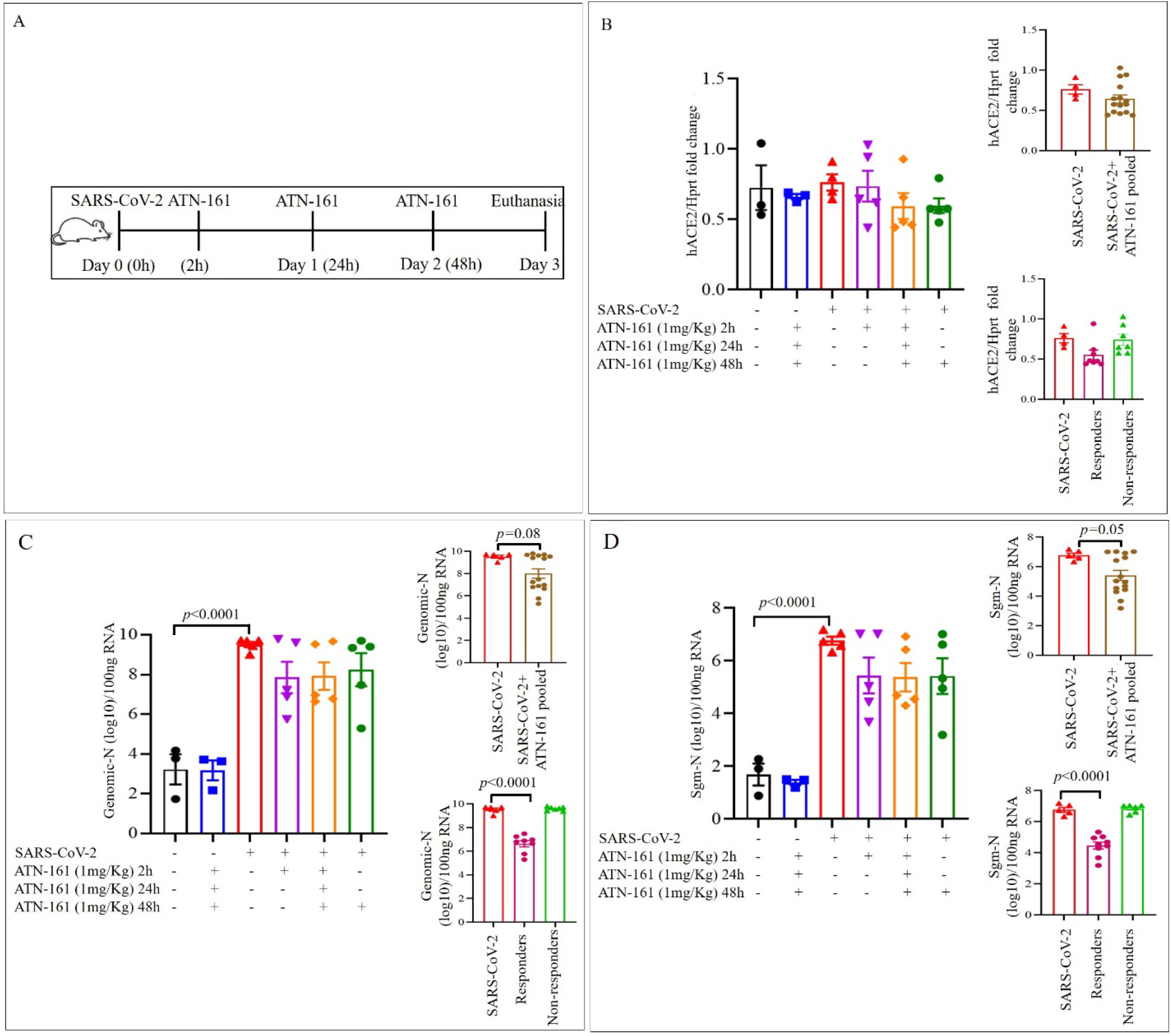
ATN-161 reduces the Genomic-N and Sub genomic N (Sgm-N) viral load in SARS-CoV-2 infected K18-hACE2 mice lung tissue. Schematic overview of experimental timeline for K18-hACE2 mice (A) 10-week old male K18-hACE2 transgenic mice administered saline (black bar) or ATN-161 (1 mg/kg, blue bar) intravenously (via retro-orbital). Mice were inoculated via the intranasal route with Severe acute respiratory syndrome coronavirus 2 (SARS-CoV-2) (2×10^5^ TCID50) + Saline (i.v. by retroorbital rout) (Red bar). ATN-161 (1 mg/Kg) treatment was administered at 3 different time periods, SARS-CoV-2+ATN-161-2h post infection (purple bar), SARS-CoV-2+ATN-161-daily (2 h, 24 h and 48 h) administration (orange bar), and SARS-CoV-2+ATN-161-48 h administration (green bar) post SARS-CoV-2 intranasal inoculation. Figure B, C, and D top inserts with brown bar represents SARS-CoV-2+ATN-161 all treatment groups pooled data; figure B, C, and D bottom insert with pink bar represents responding mice and light green bar represents non responding with SARS-CoV-2+ATN-161 (either 2, daily or 48h) treatment. 3 days post infection (3 dpi) the mice were euthanized, and RNA isolated from the left lungs by Trizol method for qRT-PCR (B) hACE2 expression in lungs of K18-hACE2 mice (C) viral genomic-N (Total-N) and (D) sub genomic-N mRNA (sgm-N). Experimental groups are divided into 6 groups. saline n=3; ATN-161 n=3; SARS-CoV-2 (5 mice for vehicle, 5 mice for each 3 ATN-161 groups). Data are presented as mean ± SEM. P values represent saline vs SARS-CoV-2+vehicle and SARS-CoV-2+vehicle vs SARS-CoV-2 +ATN-161 treatment groups.

For Genomic-N (Fig 1C), the omnibus one-way ANOVA was significant [*F*(5,20)=12.84, *p*<0.0001]. We followed up this significant analysis with two group planned comparisons using the Holm-Sidak correction. We found a significant increase in Genomic-N among SARS-CoV-2 + Saline mice compared to non-infected, Saline-treated mice [*t*(6)=5.98, *p*<0.0001]. Though we did not detect significant group differences between any other two-group comparison, visual inspection of the graph revealed that there was 1) heterogeneity among the ATN-161 treated groups, suggesting a dichotomy in this population with regards to response to the ATN-161 treatment with regard to the viral load and 2) a general trend towards reduced viral load among all ATN-161 treated groups, regardless of timepoint or number of injections. This indicated to us a potential for our analyses to be underpowered. Therefore, to increase power and reduce variability, we pooled all ATN-161 treated animals into a single group and re-analyzed the data by comparing lung viral load in this group to that of the SARS-CoV-2 + Saline-treated mice. We found a trend towards significance [*t*(18)=2.03, *p*<0.06] such that ATN-161-treated mice had lower Genomic N viral load in lungs than SARS-CoV-2-infected mice. Then we analyzed the data using the Mann-Whitney U nonparametric test to account for a significant (*p*<0.01) violation of the homogeneity of variance assumption (though ANOVA is robust to this type of violation), the effect remained marginal (*p*=0.08). We also dichotomized ATN-161-treated animals into ‘responder’ and ‘non-responder’ groups such that mice were considered non-responders if they displayed Genomic N values > 2×10^9^ (one power of 10 lower than the lowest value observed in the SARS-CoV-2 group), Sgm-N values > 1×10^5^ (one power of 10 lower than the lowest value observed in the SARS-CoV-2 group), and viral immunohistology staining counts >0.7 (one power of 10 lower than the lowest value observed in the SARS-CoV-2 group). Ns of responders for each ATN-161 treated group were 3, 3, and 2 for ATN-161-2hr, ATN-161-daily, and ATN-161-48hr groups respectively. When we re-analyzed the data by comparing lung viral load in these groups to that of the SARS-CoV-2 + Saline-treated mice, we found a significant omnibus one-way ANOVA [*F*(2,17)=35.32, *p*<0.0001] such that ATN-161 responders had significantly lower genomic lung viral loads than SARS-CoV-2-infected animals [*t*(17)=7.04, *p*<0.0001].

For Sgm-N (Fig 1D), the omnibus one-way ANOVA was significant [*F*(5,20)=13.81, *p*<0.0001] such that there was a significant increase in Sgm-N among SARS-CoV-2 + Saline mice compared to non-infected, Saline-treated mice [*t*(6)=6.13, *p*<0.0001]. Analysis of Sgm-N values among pooled ATN-161-treated mice again revealed that ATN-161 reduced Sgm-N viral load [*t*(18)=2.23, *p*<0.05] that persisted (*p*=0.05) when the Mann-Whitney U non-parametric analysis was applied given the significant (*p*<0.05) homogeneity of variance in this group. When we re-analyzed the data by comparing lung viral load in the responder/non-responder groups to that of the SARS-CoV-2 + Saline-treated mice, we found a significant omnibus one-way ANOVA [*F*(2,17)=39.91, *p*<0.0001] such that ATN-161 responders had significantly lower Sgm-N lung viral loads than SARS-CoV-2-infected animals [*t*(17)=7.40, *p*<0.0001].

### ATN-161 effects on the expression of virus in SARS-CoV-2 infected K18-hACE2 mice lung tissue

Given the observations in Figure 1, we continued our analyses comparing non-infected Saline or ATN-161-treated mice with SARS-CoV-2+Saline and either pooled ATN-161-treated infected mice or ATN-161 responders/non-responders. Immunohistochemistry staining for SARS-CoV-2 positive cells was conducted on all mice. Representative SARS-CoV-2 viral staining images from saline/ATN-161 (i); SARS-CoV-2+vehicle (ii); Responders (iii) and Non-responders (iv) with SARS-CoV-2+ATN-161 administration either 2, daily or 48 h administration, post SARS-CoV-2 intranasal inoculation are depicted in Fig 2A. All K-18 hACE2 mice infected with SARS-CoV-2 + Saline or non-responders from SARS-CoV-2+ATN-161 administration group had multifocal regions of SARS-CoV-2–positive cells (ii, iv), whereas treatment with ATN-161, 2 h and daily administered groups show 3 mice are completely negative for SARS-CoV-2 protein and 2 mice are negative (responders) in 48h ATN-161 group out of 5 mice challenged for SARS-CoV-2.

**Figure 2:**
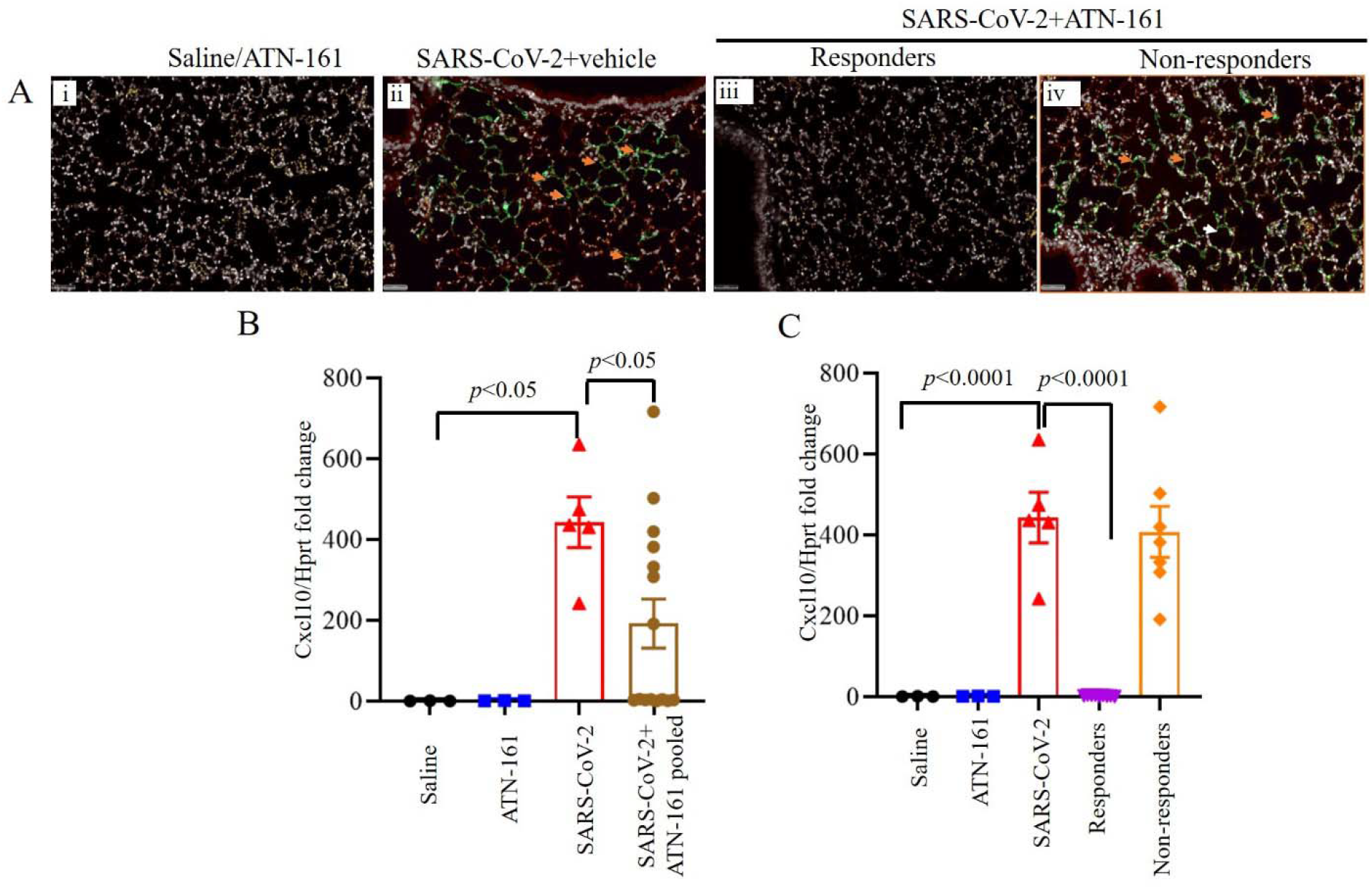
ATN-161 reduces the expression of virus in SARS-CoV-2 infected K18-hACE2 mice lung tissue. Immunohistochemistry staining for SARS-CoV-2 (Ai-iv, orange arrowheads indicates SARS-CoV-2 viral stain (green) positive cells). SARS-CoV-2 viral staining representative image from saline/ATN-161 (i); SARS-CoV-2+vehicle (ii); Responders (iii) and Non-responders (iv) with SARS-CoV-2+ATN-161 administration either 2, daily or 48 h administration, post SARS-CoV-2 intranasal inoculation. The tissue analysis at 3 dpi using the protocol described in Figure 1. All K-18 hACE2 mice infected with SARS-CoV-2 + Saline or non-responders from SARS-CoV-2+ATN-161 administration group with have multifocal regions of SARS-CoV-2–positive cells (ii, iv), whereas treatment with ATN-161, 2h and daily administered groups show 3 mice are completely negative for SARS-CoV-2 protein and 2 mice are negative (responders) in 48h ATN-161 group out of 5 mice challenged for SARS-CoV-2. (B-C) Cxcl10 mRNA expression in lung. Scale bars, 50 µm. Green = SARS-CoV-2; White = nuclei/DAPI; Red = empty/autofluorescence. Data are presented as mean ± SEM. P values represent saline vs SARS-CoV-2+vehicle and SARS-CoV-2+vehicle vs SARS-CoV-2+ATN-161 pooled, SARS-CoV-2+ATN-161 responders, and SARS-CoV-2+ATN-161 non-responders.

One-way ANOVA of C–X–C motif chemokine ligand 10 (*Cxcl10*) mRNA expression in lung comparing saline or ATN-161 treated non-infected, SARS-CoV-2 + Saline, and SARS-CoV-2 + ATN-161 pooled mice (Fig 2B) revealed a significant effect [*F*(3,23)=4.56, *p*<0.05] such that SARS-CoV-2+Saline mice had significantly higher lung *Cxcl10* mRNA expression compared to Saline-treated uninfected mice [*t*(6)=3.07, *p*<0.05] and pooled ATN-161-treated mice [*t*(18)=2.46, *p*<0.05]. One-way ANOVA of *Cxcl10* mRNA expression in lung comparing saline or ATN-161 treated non-infected, SARS-CoV-2 + Saline, and SARS-CoV-2 + ATN-161 responders and non-responders (Fig 2C) revealed a significant effect [*F*(4,21)=24.38, *p*<0.0001], that persisted when the Kruskal-Wallis non-parametric analysis (*p*<0.005) was applied given the significant violation of homogeneity of variance (*p*<0.05) in this ANOVA analysis. Two-group planned follow-up comparisons revealed that SARS-CoV-2+Saline mice had significantly higher lung *Cxcl10* mRNA expression compared to Saline-treated uninfected mice [*t*(6)=5.59, *p*<0.0001] and ATN-161-treated responders [*t*(11)=7.12, *p*<0.0001].

### Histopathological Analysis of SARS-CoV-2 infected K18-hACE2 mice lung tissue by Hematoxylin and eosin stain

Representative images of hematoxylin and eosin staining of lung sections from saline/ATN-161 (Fig. 3Ai-Aii); SARS-CoV-2+vehicle (Fig. 3Bi-ii); Responders (Fig. 3Ci-ii) and non-responders (Fig. 3Di-ii) with SARS-CoV-2+ATN-161 (either 2h, daily, or 48h) administration post-SARS-CoV-2 intranasal inoculation are shown in Figure 3. Histopathological observations indicated that multifocal lesion, moderate interstitial pneumonia (Fig. 3Bii, Dii, black frames), infiltration of lymphocytes (Fig. 3Bii, Dii, green arrow), fibrin exudation (Fig. 3Bii, black arrow) were observed in SARS-CoV-2+vehicle and SARS-CoV-2+ATN-161 non responders mice whereas these observations are less or completely absent in SARS-CoV-2+ATN-161 responders mice. Hematoxylin and eosin staining of lung sections from saline/ATN-161 (Fig. 3Ai-Aii) revealed no histopathological changes in any of the mice.

**Figure 3.**
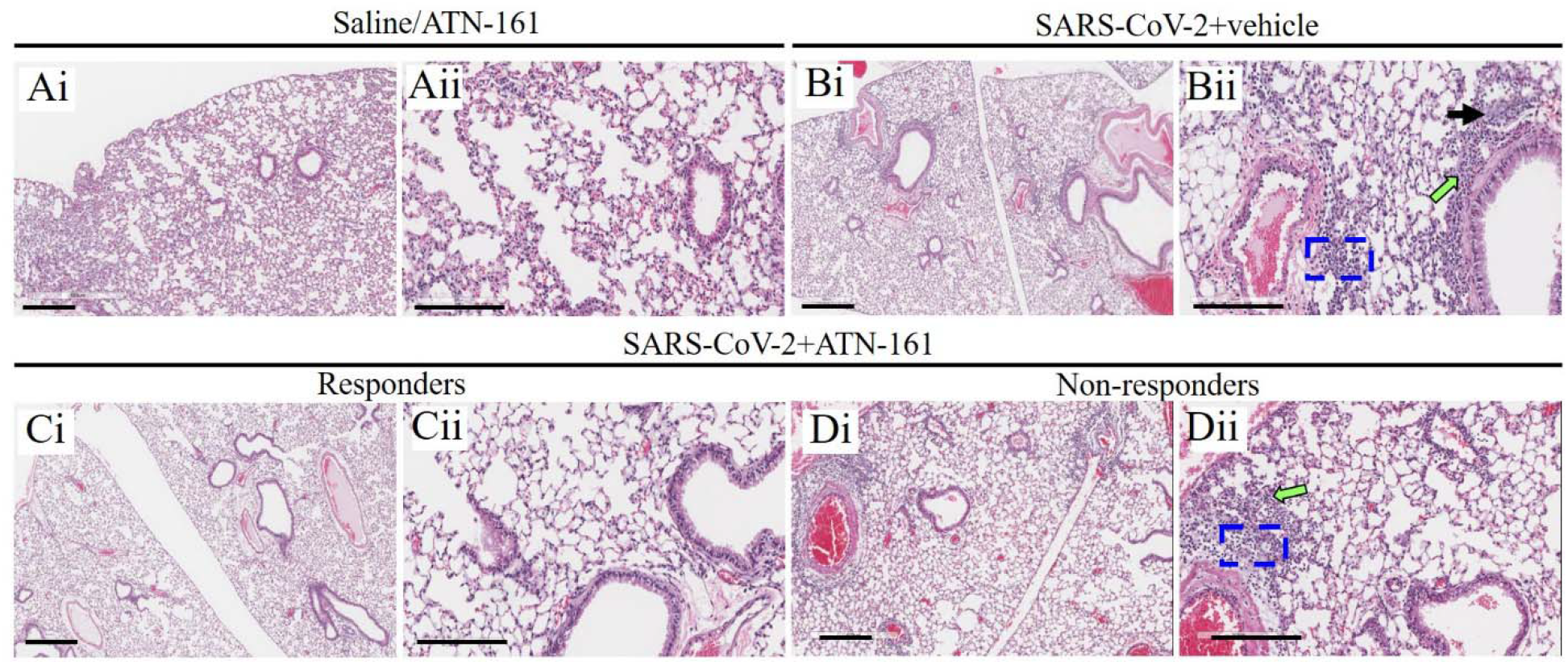
Histopathological Analysis of SARS-CoV-2 infected K18-hACE2 mice lung tissue. (Ai–Dii) Hematoxylin and eosin staining of lung sections from saline/ATN-161 (Ai-Aii); SARS-CoV-2+vehicle (Bi-ii); Responders (Ci-ii) and Non-responders (Di-ii) with SARS-CoV-2+ATN-161(either 2h, daily, or 48h) administration post SARS-CoV-2 intranasal inoculation with tissue analysis at 3 dpi using the protocol described in Figure 1. Histopathological observations indicated that multifocal lesion, moderate interstitial pneumonia, (Bii, Dii, black frames), infiltration of lymphocytes (Bii, Dii, green arrow), fibrin exudation (Bii, black arrow). Each image is representative of a group of Control n=3; ATN-161 n=3; SARS-CoV-2 (5 mice for vehicle, 5 mice for each 3 ATN-161 groups). Scale bar: 500 μm (Ai, Bi, Ci, and Di) and 200 μm (Aii, Bii, Cii, and Dii).

### Histopathological Analysis of SARS-CoV-2 infected K18-hACE2 mice lung tissue Masson’s Trichrome stain

Representative images of Masson’s Trichrome-stained sections (Fig. 4Ai–Dii) of lung sections from saline/ATN-161 (Fig. 4Ai-Aii); SARS-CoV-2+vehicle (Fig. 4Bi-ii); Responders (Fig. 4Ci-ii) and Non-responders (Fig. 4Di-ii) with SARS-CoV-2+ATN-161 (either 2h, daily, or 48h) administration post SARS-CoV-2 intranasal inoculation. Histopathological analysis revealed multiple intra-arteriolar microthrombi (black arrows), intra-alveolar microthrombi (green arrows), large interstitial hemorrhagic area (yellow arrow) in SARS-CoV-2+vehicle and SARS-CoV-2+ATN-161 non responders mice whereas these observations are less or completely absent in SARS-CoV-2+ATN-161 responders mice. Masson’s Trichrome staining of lung sections from saline/ATN-161 (Fig. 4Ai-Aii) were no histopathological changes in any of the mice.

**Figure 4.**
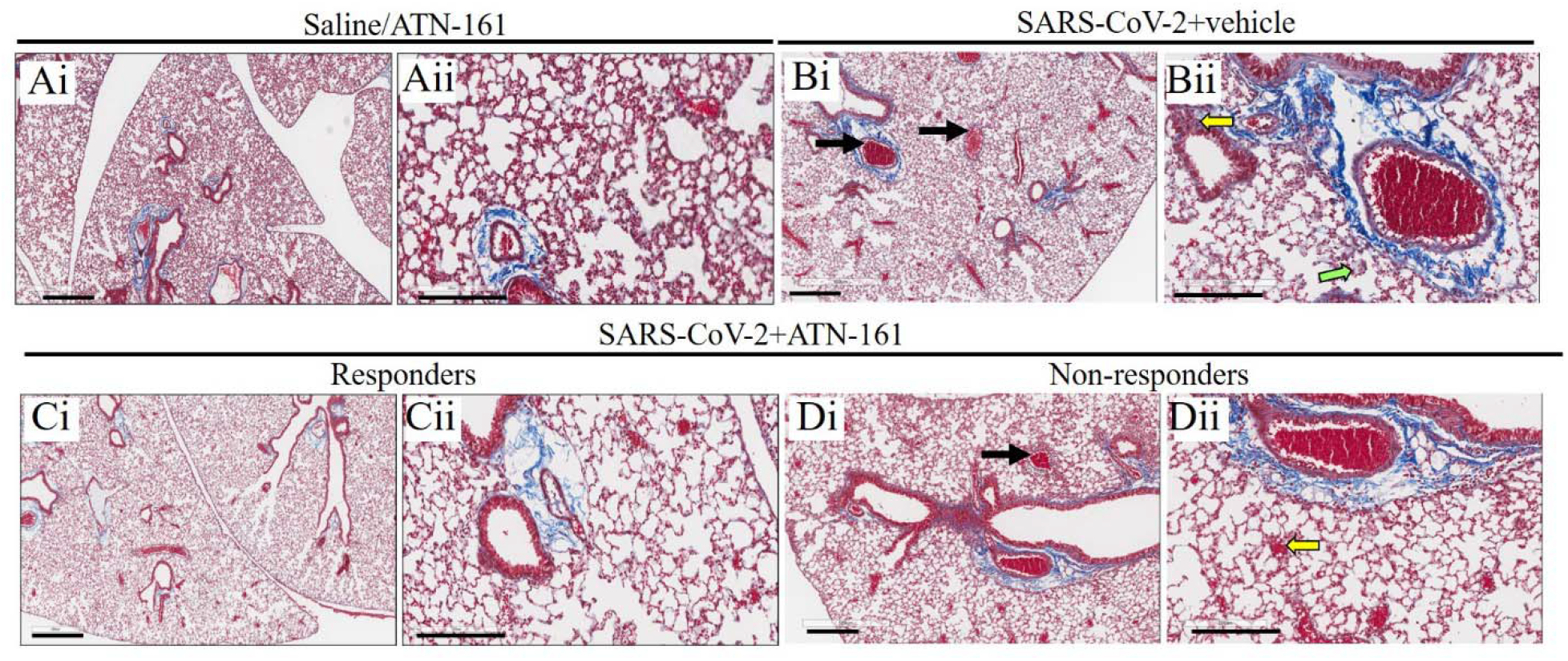
Histopathological Analysis of SARS-CoV-2 infected K18-hACE2 mice lung tissue. Masson’s Trichrome-stained sections (Ai–Dii) of lung sections from Saline/ATN-161 (Ai-Aii); SARS-CoV-2+vehicle (Bi-ii); Responders (Ci-ii) and Non-responders (Di-ii) with SARS-CoV-2+ATN-161(either 2h, daily, or 48h) administration post SARS-CoV-2 intranasal inoculation with tissue analysis at 3 dpi using the protocol described in Figure 1. Histopathological observations indicated that showing multiple intra-arteriolar microthrombi (black arrows), intra-alveolar microthrombi (green arrows), large interstitial hemorrhagic area (yellow arrow). Each image is representative of a group of Control n=3; ATN-161 n=3; SARS-CoV-2 (5 mice for vehicle, 5 mice for each 3 ATN-161 groups). Scale bar: 500 μm (Ai, Bi, Ci, and Di) and 200 μm (Aii, Bii, Cii, and Dii).

**Figure 5.**
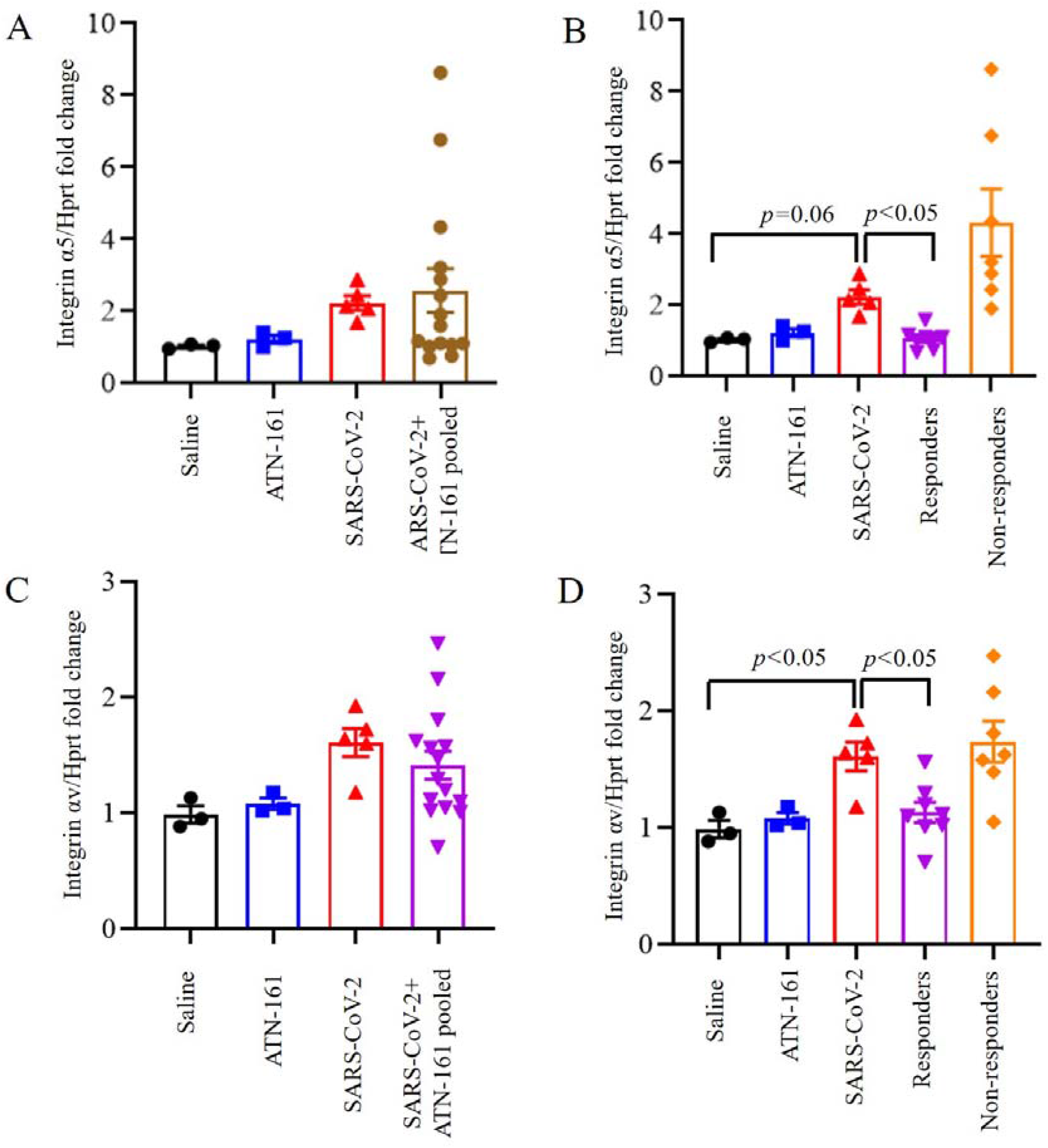
Induced expression of integrin α5, and integrin αv in the lungs of SARS-CoV-2 infected K18-hACE2 mice and is inhibited by ATN-161 treatment. Integrin α5 (A-B), and Integrin αv (C-D) mRNA expression in lung at different time periods of ATN-161 by intravenous injection (i.v) after infection with SARS-CoV-2 (TCID50= 2×10^5^ /0.05 mL) was determined by qRT-PCR. Data are presented as mean ± SEM. P values represent saline vs SARS-CoV-2+vehicle and SARS-CoV-2+vehicle vs SARS-CoV-2+ATN-161 pooled, SARS-CoV-2+ATN-161 responders, and SARS-CoV-2+ATN-161 non-responders.

### Induced expression of integrin α5, and integrin αv in the lungs of SARS-CoV-2 infected K18-hACE2 mice and is inhibited by ATN-161 treatment

One-way ANOVA of integrin-α5 expression in lung comparing saline or ATN-treated non-infected, SARS-CoV-2 + Saline, and SARS-CoV-2 + ATN-161 pooled mice was not significant (Fig 4A). One-way ANOVA of integrin-α5 expression in lung comparing saline or ATN-161 treated non-infected, SARS-CoV-2 + Saline, and SARS-CoV-2 + ATN-161 responders and non-responders (Fig 4B) revealed a significant effect [*F*(4,21)=6.59, *p*<0.05], that persisted when the Kruskal-Wallis non-parametric analysis (*p*<0.05) was applied given the trend towards a violation of homogeneity of variance in this ANOVA analysis (*p*=0.06). Because there was very high variability in the SARS-CoV-2 + ATN-161 non-responders, and we were concerned about the possibility of inflated Type 1 error risk, we conservatively conducted two-group follow-up comparisons using the non-parametric approach (Dunn-corrected). We found that SARS-CoV-2+Saline mice tended to have higher lung integrin-α5 expression compared to uninfected Saline-treated mice [*z*=2.32, *p*=0.06].

One-way ANOVA of integrin-αv expression in lung comparing saline or ATN-161 treated non-infected, SARS-CoV-2 + Saline, and SARS-CoV-2 + ATN-161 pooled mice was not significant (Fig 4C). One-way ANOVA of integrin-αv expression in lung comparing saline or ATN-161 treated non-infected, SARS-CoV-2 + Saline, and SARS-CoV-2 + ATN-161 responders and non-responders (Fig 4D) revealed a significant effect [*F*(4,21)=6.04, *p*<0.005], such that SARS-CoV-2+Saline mice had significantly higher lung integrin-αv expression compared to Saline-treated uninfected mice [*t*(6)=2.70, *p*<0.05] and ATN-161-treated responders [*t*(11)=2.67, *p*<0.05].

## Discussion

The coronavirus disease-2019 (COVID-19) caused by SARS-CoV-2 continues to ravage the world. As of March 24, 2021, there were 3,089,162 deaths with a total of 145,759,060 confirmed (COVID-19) cases worldwide [38, 39]. While the development of vaccines and antibody treatments against COVID-19 is promising and globally accepted in the current pandemic [40], the emergence of SARS-CoV-2 viral variants may present major challenges to these therapeutic approaches. Like most viruses, SARS-CoV-2 mutates and continually presents variants. Spike protein mutations are the most common mutation seen in SARS-CoV-2 variants, which facilitate viral entry into the cell and mediate viral propagation by binding to ACE2 receptor. Interestingly, an RDG integrin-binding motif is a novel feature of SARS-CoV-2 spike protein, which is not seen in other coronaviruses [41, 11]. While this feature may have enhanced viral infectivity of SARS-CoV-2, it remains unknown how variant strains of SARS-CoV-2 may affect binding to integrins, with the resultant feature of affecting viral entry and propagation. However, it is worth noting that to the best of our knowledge, none of the currently characterized SARS-CoV-2 variants have <u>directly</u> mutated the RGD motif; This suggests the possibility that the RGD motif is of evolutionary advantage to the virus by supporting its ability to infect hosts.

Integrins are cell surface receptors that may bind to the SARS-CoV-2 spike protein interaction [42, 43, 11, 33]. In our previous study, we demonstrated by ELISA that the SARS-CoV-2 spike protein and ACE2 could bind to α5β1, and that ATN-161 could disrupt both of these interactions [26]. We further demonstrated that ATN-161 could significantly reduce SARS-CoV-2 infection (viral load, cell viability and cytopathy) in Vero (E6) cells in vitro [26]. In this study, we extend the understanding of the role of integrins and explored the therapeutic role of ATN-161 against SARS-CoV-2 infection in vivo in the k18-hACE2 mice model.

K18-hACE2 mice provide a platform for the rigorous screening of candidate drugs before their evaluation in other animal models [44]. This mouse model is widely used to evaluate the pathogenicity of viruses such as SARS-CoV-2 that require or prefer the human form of ACE2 (versus mouse ACE2) to readily infect mice and can be used to study potential therapies [45, 35]. Previous studies provided the evidence that SARS-CoV-2 infection could cause typical interstitial pneumonia and develop respiratory disease in hACE2-expressing mice resembling what is commonly seen in COVID-19 patients [46, 45, 47, 48].

We measured genomic-N (Nucleocapsid (N) protein) and Sgm-N in lungs to assess SARS-CoV-2viral infectivity and replication. Our results showed that SARS-CoV-2–infected hACE2 mice had significantly higher SARS-CoV-2 genomic-N. Similarly, viral sgmRNA copies were detected predominantly in the lung as compared to uninfected saline treated mice. SgmRNA levels of the virus is an adequate surrogate assay for detection of replicating virus (replicating virus separated from the total genome, and form dimers as the virus is replicating its machinery which can continue to produce protein) [34]. Thus, we are confident that our mice were successfully infected with SARS-CoV-2 [35].

We found that ATN-161-treated mice had lower Genomic-N viral load in lungs than saline treated SARS-CoV-2-infected mice. Visual inspection of Genomic-N and sgm-N graph revealed that there was 1) heterogeneity among the ATN-161 treated groups, suggesting a dichotomy in this population with regards to ATN-161 treatment response and viral load, and 2) a general trend towards reduced viral load among all ATN-161 treated groups, regardless of timepoint or number of injections. A limitation of this study is that it employed a limited number of K18 hACE2 mice, a function of the logistical difficulty of doing live virus BSL-3 studies. We also dichotomized ATN-161-treated animals into ‘responder’ and ‘non-responder’ groups such that mice were considered non-responders if they displayed Genomic N values > 2×10^9^, Sgm-N values > 1×10^5^ and viral immunohistology staining counts >0.7 values observed in the SARS-CoV-2 group in the SARS-CoV-2 + saline group. ATN-161 responders had significantly lower genomic lung viral loads than SARS-CoV-2-infected animals.

Accordingly, this bimodal distribution of responders may be due to our use of heterozygous (HT) K18-hACE2 mice, as the K18-hACE2 homozygous mouse model completely replaces mACE2 expression with hACE2 under the mAce2 promoter [49]. Thus, it might be possible that our use of HT K18-hACE2 mice in this study produced variable expression patterns of mACE2 vs hACE2, reducing the efficacy of ATN-161 in mice that expressed higher levels of mACE2 relative to hACE2. ATN-161 interference with mACE2-α5β1 interactions would presumably have minimal to no effect on SARS-CoV-2 infection as compared to impacting hACE2-α5β1 interactions. ATN-161 reduced Sgm-N viral load among pooled ATN-161-treated mice. When we re-analyzed the data by comparing lung viral load in the responder/non-responder groups to that of the SARS-CoV-2 + Saline-treated mice, we found that ATN-161 responders had significantly lower Sgm-N lung viral loads than SARS-CoV-2-infected animals. This suggests a lack of statistical power due to low Ns among individual ATN-treated groups; follow-up studies with additional subjects will be conducted to replicate these findings.

All K-18 hACE2 mice infected with SARS-CoV-2 or non-responders from SARS-CoV-2+ATN-161 administered group had multifocal regions of SARS-CoV-2–positive cells. Our results are further supported by recent findings of lung histological changes in SARS-CoV-2 infected hACE2 mice [35, 49]. Furthermore, we observed that viral immunohistology staining counts were negative in responders for each ATN-161 treated group with n’s of 3, 3, and 2 for ATN-161-2 h, ATN-161-daily, and ATN-161-48 h groups, respectively. The infected K18-hACE2 mice did not lose body weight after only 3dpi as expected, but we observed high levels of viral copies and infectious virus in the lungs given as demonstrated by other studies in K18-hACE2 mice [35, 49, 50].

In regards to an inflammatory response to SARS-CoV-2 infection, we observed that *Cxcl10* mRNA expression was significantly upregulated in the lungs of SARS-CoV-2 infected mice as has previously been reported [50]. In patients having rapid early viral replication, this is followed by inflammatory responses that contribute to pathology [51]. Post-mortem analysis of human COVID-19 patients showed immune cell accumulation in the lungs [52]. In our study, ATN-161 treatment, either pooled or ATN-161 responders, significantly lowered SARS-CoV-2-induced *Cxcl10* levels, a particularly robust result suggesting that ATN-161 has anti-inflammatory properties in the context of SARS-CoV-2 infection. This result is in agreement with our previous studies in experimental ischemic stroke where post-stroke treatment with ATN-161 reduced neuro-inflammation [31].

Morphological changes observed at 3 dpi in infected lungs of K18-hACE2 mice included multifocal lesions, moderate interstitial pneumonia, infiltration of lymphocytes, and fibrin exudation. These were seen in SARS-CoV-2+vehicle and SARS-CoV-2+ATN-161 non responders mice but were less or completely absent in SARS-CoV-2+ATN-161 responder mice as well as absent in uninfected saline/ATN-161 treated mice. These findings are consistent with previous reports from post-mortem examination of patients with COVID-19 and SARS-CoV-2-infected K18-hACE2 mice in other reports [50, 49, 53-55].

Pulmonary fibrosis, a particularly negative consequence of COVID-19-induced acute respiratory distress syndrome [56], was not prominentin infiltrated areas at 3 dpi in infected lungs of K18-hACE2 mice. However, we observed moderate multiple intra-arteriolar microthrombi, intra-alveolar microthrombi, and large interstitial hemorrhagic areas in SARS-CoV-2+vehicle and SARS-CoV-2+ATN-161 non responder mice whereas these observations were less or completely absent in SARS-CoV-2+ATN-161 responder mice. Severely infected COVID-19 patient autopsy samples showed that microthrombi or immunothrombi is associated with a hyperinflammatory response [56, 57]. Our findings are further supported by recent results in the human immune system humanized mouse model on histological changes in the lungs with SARS-CoV-2 infection [58, 35, 49].

Our results suggest a critical role of integrins as an additional receptor to SARS-CoV-2 spike protein cell entry [59, 60]. We observed induced expression of integrin α5 and integrin αv in the lungs of SARS-CoV-2 infected K18-hACE2 mice whereas lung expression of hACE2 levels did not vary in SARS-CoV-2+Saline or ATN-161 treated mice, suggesting that SARS-CoV-2 infection and / or pathogenesis involves these, and perhaps other integrins, that activate downstream signaling to induce lung pathology [33, 61, 62]. Indeed, a recent study showed that increased integrin α5β1 and αvβ3 levels in cardiac myocytes, obtained from heart failure patients, correlates with ACE2 expression [63]. This suggests that the concomitant elevation of these integrins and the upregulation of ACE2 in an organ may render it more susceptible to SARS-CoV-2 infection. Hence, as observed in the present study, decreasing the expressions of α5β1 and αVβ3 in the presence of ACE2 may dampen the effect of SARS-CoV-2 infection on lung tissue morphology. Other studies have clearly linked integrin α5β1, and its inhibition, with other viral infections [62, 64]. ATN-161 treatment inhibited porcine hemagglutinating encephalomyelitis virus (PHEV) infection and its increase in the expression of integrin α5β1 *in vivo* in a mouse model [64]. Similarly, we observed that ATN-161 treatment inhibits SARS-CoV-2-induced integrin α5, and integrin αv in the K18-hACE2 mice lungs among ATN-responders. Importantly, our studies were performed at the dose of ATN-161, 1 mg/kg, that demonstrated efficacy in *in vitro* and *in vivo* preclinical studies in blocking angiogenesis, solid tumor growth, and ischemic stroke injury [65, 30, 31]. This dose was also similar to the 0.8 mg/kg dose used in the PHEV anti-viral *in vivo* studies [64]. However, as we and others have demonstrated U-shaped dose response effects with ATN-161 [30, 26] additional *in vivo* SARS-CoV-2 dosing studies are necessary and currently underway. However, ATN-161 present several potential advantages as a novel COVID-19 therapy. Unlike other RGD-based peptides, it preferentially binds to and inhibits <u>activated</u> forms of α5β1 in areas of inflammation, hypoxia and angiogenesis [30]. ATN-161 also binds to integrin αvβ3 [30], an additional integrin that has been implicated in SARS-CoV-2 pathogenesis [33, 32], and is safe, well-tolerated in human clinical trials (cancer) with no dose limiting toxicity [28, 66], and can be administered i.v., i.p., and intranasally [31, 67]; The latter may support a more readily accessible means of COVID-19 treatment as well as afford a prophylactic approach which is currently under investigation in our laboratory.

## Conclusion

To the best of our knowledge, this study is the first to demonstrate that integrin blockade can reduce SARS-CoV-2 infection *in vivo*. Specifically, we demonstrated that post-infection treatment with ATN-161 reduces lung viral load, replication, improves lung histology and reduces lung α5 and αv integrin expression in a majority of SARS-CoV-2 infected k-18 mice. Our results further support the hypothesis that integrins play an important role in SARS-CoV-2 infection as well as support the further investigation of ATN-161, and potentially other anti-integrin agents, as novel therapies for COVID-19.

### Future perspective

Although several studies have predicted the potential for SARS-CoV-2 to bind integrins and thereby infect host cells with or without associated ACE2 interaction, for the first time we present evidence here that inhibition of integrin α5β1 (and αvβ3) *in vivo* can reduce both SARS-CoV-2 viral load and pathological complications in lung tissue of SARS-CoV-2 in k18-hACE2 mice. Despite the limitations of our study (small sample size, use of heterozygous k18-hACE2 mice), the promise for ATN-161 as a potential therapeutic and prophylaxis after SARS-CoV-2 exposure remains attractive. To date, several small-molecule drug and peptide treatments have been developed for intranasal delivery which reach systemic circulation rapidly, many of which have been formulated to treat diseases which have the ability to alter cognition, such as depression [68]. As the olfactory and trigeminal nerves offer a safe and effective delivery pathway to deliver therapeutic agents to the brain, an intranasal-based therapy to both prevent and treat neurological complications of post-acute COVID-19 syndrome may one day include a formulation of ATN-161 [69, 70].

## AUTHOR CONTRIBUTIONS

N.A performed experiments, N.A, E.B.E-C, designed, analyzed the data, and wrote the manuscript. I.B.M., W.C. assisted in performing experiments. T.G., BIJ, performed literature searches and drafted discussion, and G.J.B. conceived, designed and supervised the experiments, and revised and finalized the manuscript. All authors have read and approved the final manuscript.

## Acknowledgements

We would like to thank Drs. Jay K. Kolls, Xuebin Qin (Tulane University) for their thoughtful comments and advice on this manuscript. Angela Birnbaum, Kristin E. Chandler, Thompson, Carli C for BSL3 service (Tulane University). Authors are grateful to Naoki Iwanaga, Smither, Allison R, Brandon J. Beddingfield, Nicholas J. Maness for technical help in experiments. Authors are thankful to Cecily C. Midkiff, Confocal Microscopy and Molecular Pathology Core, Dina Guapp, Histology Core, Delucca, Beatris, Tulane University. The authors appreciate Rebecca Solch, Meshi Paz, Jaime Befeler, Hawkins, Scott V, Department of Neurosurgery at Tulane University, for assistance.

## Funding

Tulane University startup funds. to G.J.B and NIH/NIMH K01 MH117343 to E.B.E-C.

## Conflicts of Interest

The authors have declared that no conflict of interest exists.

## Notes

### Competing Interest Statement

The authors have declared no competing interest.

